# Estradiol and luteinizing hormone reverse memory loss in phencyclidine model of schizophrenia: Evidence for hippocampal GABA action

**DOI:** 10.1101/207159

**Authors:** Alexander J. Riordan, Ari W. Schaler, Jackson B. Fried, Tracie A. Paine, Janice E. Thornton

**Author notes:** **Corresponding author:** Alexander J. Riordan, Princeton Neuroscience Institute, Princeton University, Washington Road, Princeton, NJ 08544, Tel: N/A, Fax: N/A.

## Abstract

The cognitive symptoms of schizophrenia are poorly understood and difficult to treat. Estrogens may mitigate these symptoms via unknown mechanisms. To examine these mechanisms, we tested whether increasing estradiol (E) or decreasing luteinizing hormone (LH) could rescue declarative memory in a phencyclidine (PCP) model of schizophrenia. We then assessed whether changes in cortical or hippocampal GABA may underlie these effects. Female rats were ovariectomized and injected subchronically with PCP. To modulate E and LH, animals received hormone capsules or Antide injections. Short-term episodic memory was assessed using the novel object recognition task. Brain expression of GAD67 was analyzed via western blot, and parvalbumin-containing cells were counted using immunohistochemistry. Some rats received hippocampal infusions of a GABA_A_ agonist, GABA_A_ antagonist, or GAD inhibitor before behavioral testing. We found that PCP reduced hippocampal GAD67 and abolished object recognition. Antide restored hippocampal GAD67 and rescued recognition memory in PCP-treated animals. Estradiol reversed PCP’s amnesic effect but failed to restore hippocampal GAD67. PCP did not cause significant differences in number of parvalbumin-expressing cells or cortical expression of GAD67. Hippocampal infusions of a GABA_A_ agonist restored memory in PCP-treated rats. Blocking hippocampal GAD or GABA_A_ receptors in ovx animals reproduced memory loss similar to PCP and inhibited estradiol’s memory rescue in PCP-treated animals. In summary, decreasing LH or increasing E can reverse memory loss in a PCP model of schizophrenia. Alterations in hippocampal GABA may contribute to both PCP’s effects on declarative memory and the hormones’ ability to reverse them.

## 1. INTRODUCTION

Schizophrenia is a common neuropsychiatric disorder that causes devastating cognitive, negative, and positive symptoms (McGrath et al. 2008). Despite the importance of cognitive symptoms for a patient’s long-term well-being, current treatments for schizophrenia fail to adequately improve them (Green et al. 2004). Better therapies and a greater understanding of the brain mechanisms underlying cognitive loss in schizophrenia are needed.

A dominant model of schizophrenia suggests that hypofunctioning N-methyl-d-aspartate glutamate receptors (NMDARs) play a role in generating schizophrenia symptoms (Moghaddam and Javitt 2012). Repeated exposure to NMDAR antagonists causes schizophrenia-like symptoms in healthy humans and exacerbates symptoms in those with schizophrenia (Jones et al. 2011; Moghaddam and Javitt 2012; Nakazawa et al. 2012). Based on these findings, animal models of schizophrenia using repeated (sub-chronic) treatment with NMDAR antagonists such as phencydlidine (PCP) and ketamine have been developed (Jones et al. 2011; Moghaddam and Javitt 2012). Animals in a number of species that have been subjected to these regimens show traits that approximate positive, negative, and cognitive symptoms of schizophrenia, including loss of short-term episodic memory (Jones et al. 2011; Rajagopal et al. 2014). Although no animal model perfectly replicates schizophrenia, the unique cross-species validity of NMDAR antagonist models has rendered them enduring tools for studying possible mechanisms and treatments for the disorder (Jones et al. 2011).

Current theories propose that NMDA hypofunction disrupts gamma-amino-butyric-acid (GABA) neurons, causing cognitive loss in schizophrenia (Lewis et al. 2012). Experiments suggest that abnormal GABA-containing inhibitory neurons are common in schizophrenia (Nakazawa et al. 2012). Post-mortem analyses reveal that the brains of people with schizophrenia express lower amounts of the rate-limiting enzyme for GABA synthesis, glutamic acid decarboxylase 67 (GAD67) (Benes et al. 2007; Lewis et al. 2012). There is also reduced expression of calcium-binding protein parvalbumin (PV), which marks a subset of GABAergic neurons (Beasley and Reynolds 1997; Zhang and Reynolds 2002). These GABAergic abnormalities are found in a number of brain regions thought to be important for declarative and short-term memory, including the hippocampus and frontal neocortex (Benes et al. 2007; Lewis et al. 2012; Zhang and Reynolds 2002). Studies in NMDAR animal models of schizophrenia have found similar deficits of GAD67 and PV expression in the prelimbic cortex and hippocampus, further supporting their possible mechanistic role in schizophrenia-like cognitive loss (Abdul-Monim et al. 2007; Behrens et al. 2007; Kaalund et al. 2013).

There are indications that estradiol can mitigate cognitive symptoms in schizophrenia and animal models of the disease (for review see McGregor et al. 2017). Clinical trials have suggested that estrogen and analogues may improve some measures of cognition in males and females with schizophrenia (McGregor et al. 2017). Estradiol can also improve recognition memory in the PCP model of schizophrenia when given before or after PCP treatment (Roseman et al. 2012; Sutcliffe et al. 2008). Still, the underlying mechanisms of estradiol’s cognitive rescue remain unknown, and the possibility that estradiol works through GABAergic pathways to restore memory has not been investigated.

Estrogen inhibits release of luteinizing hormone (LH) through negative feedback (Charlton 2008; Wise and Ratner 1980), and it is conceivable that estrogen’s beneficial effects in schizophrenia are due to LH inhibition. High LH levels are associated with memory decline in humans, and rodent studies support a causal relationship (Burnham et al. 2016; Burnham and Thornton 2015; Casadesus et al. 2007). Nevertheless the molecular mechanisms underlying LH’s amnesic effects remain uncharacterized, and the therapeutic possibilities of LH inhibition in schizophrenia are unknown.

Here we tested whether modulating estradiol (E) or LH could rescue recognition memory in the PCP model of schizophrenia, and if these hormones act by counteracting PCP’s putative effects on GABA neurons. To examine the actions of E and LH without the confounds of hormonal cycles, we ovariectomized female rats and systematically altered their estradiol or LH levels (Roseman et al. 2012; Wise and Ratner 1980; Ziegler and Thornton 2010). We then injected them with subchronic PCP and assessed the rats’ short-term recognition memory using the novel object recognition task (NORT) (Janhunen et al. 2015; Jones et al. 2011; Rajagopal et al. 2014; Roseman et al. 2012). NORT is widely used to assess episodic memory loss in animal models of schizophrenia, and performance in the task depends on specific functioning of cortical and hippocampal regions altered in the disorder (Benes et al. 2007; Broadbent et al. 2004; Cohen et al. 2013; Lewis et al. 2012; Oliveira et al. 2010; Rajagopal et al. 2014; Zhang and Reynolds 2002). When paired with a short intertrial interval, NORT also measures short-term memory, which is disrupted in humans with schizophrenia but improved by estrogenic compounds (McGregor et al. 2017; Rajagopal et al. 2014).

After these initial behavioral characterizations, we assessed whether the hormonal memory rescue may rely on GABAergic mechanisms. We first tested whether PCP reduced expression of GAD67 and PV in the dorsal hippocampus or prelimbic cortex and determined if our hormonal treatments restored these GABAergic markers. To assess whether changes in hippocampal GABA could account for PCP’s disruption of recognition memory and the hormones’ mnemonic rescue, respectively, we infused a GABA_A_ receptor agonist into the dorsal hippocampus of PCP-treated females and blocked hippocampal GABA_A_ receptors or GABA synthesis in non-PCP-treated controls and estradiol-treated PCP animals during NORT.

## 2. MATERIALS AND METHODS

### 2.1. Subjects

Adult female Sprague-Dawley rats (Hilltop Lab Animals, Inc., Scottdale, PA), 3-7 months old, were housed in groups of 2-3 in plastic cages (see supplementary methods). All procedures met NIH standards (National Research Council of the National Academies 2011) and were approved by the Oberlin College Institutional Animal Care and Use Committee.

### 2.2. Ovariectomy, Hormones, and PCP treatment

To control E levels, females were ovariectomized (ovx) under isoflurane anesthesia (1-3% with 1L/min O_2_) and implanted subcutaneously with either a silastic capsule filled with 17β-estradiol (E) or a blank (Blk) capsule. Because of negative feedback between E and LH, blank implants result in low E and high LH, whereas E implants of this size result in physiological levels of E and low LH (Roseman et al. 2012; Wise and Ratner 1980; Ziegler and Thornton 2010). After surgery, animals were allowed to recover for 7d.

To create an animal model of schizophrenia, females were injected bidaily, IP with either PCP (2mg/ml/kg) or vehicle (sterile 0.9% NaCl 1ml/kg) for 7d. Injections were followed by a 7d washout period, during which all animals were handled (Roseman et al. 2012).

To lower LH levels in Blk animals, Antide (Ant: 1mg/ml/kg) or vehicle (1ml/kg sterile H_2_O) was injected sub-cutaneously 6h before behavioral testing or sacrifice (Ziegler and Thornton 2010).

### 2.3. Novel Object Recognition Test (NORT) and Locomotion Test

Before testing, animals were habituated to the open field testing arena (supplementary methods). Animals then underwent novel object recognition testing as described elsewhere (Roseman et al. 2012). Object placement and type were counterbalanced to control for possible spatial or object preferences. Behavioral data were averaged for each female. Testers were blind to treatment group.

Exposure and test trial data are expressed as number of seconds animals explored each object. Test trial data are also expressed as a discrimination index *d*, where *d*=(time novel object explored – time known object explored)/(time novel object explored + time known object explored). Any test in which an animal did not explore both objects was excluded.

To assess possible drug and hormone effects on locomotion, animals were placed individually into an open field box, and the number of lines crossed during a 5min period was counted (see supplement) (Roseman et al. 2012).

### 2.4. Western Blot

Protein concentrations of GAD67 were measured in dorsal hippocampus and prelimbic cortex via western blot (Paxinos and Watson 2007) (see supplement for details). Briefly, animals were anaesthetized, decapitated, and brains were snap frozen in dry-ice-cooled isopentane (Sigma). Tissue punches from dorsal hippocampus and prelimbic cortex were sonicated, diluted to equal protein concentrations, and stored at −70°C until SDS-PAGE and transfer to nitrocellulose membranes. After blocking, membranes were incubated in mouse-anti-GAD67 antibody (Sigma, 1:2000), washed, then incubated in HRP-conjugated secondary antibody (Vector Labs, 1:5000). Membranes were exposed to chemiluminescent detection reagents (Thermo Fisher Scientific) and imaged. Membranes were stripped and re-probed for actin (Sigma, 1:10000) using the same procedure.

Densitometry was performed for all sample bands using ImageJ. To control for sample loading or transfer differences, all GAD67 values were normalized over actin levels and intra-gel Blk controls. Three technical replicates were performed for hippocampal samples, and the median value was used as the final fold value for each animal. Outliers were determined using non-biased methods described in Leys et al., 2013 (see supplement) (Leys et al. 2013).

### 2.5. Immunohistochemistry & Cell Counting

Rats were anesthetized and perfused, 50μm coronal brain sections were cut, and every fifth section was blocked then incubated in polyclonal rabbit-anti-parvalbumin antibody (Swant PV-28; 1:5000 in PBS) for 48h at 4°C. Sections were subsequently washed and incubated in biotinylated secondary antibody (goat-anti-rabbit IgG: 1:200, Vector Labs) for 45min at RT. After further washes, sections were incubated in avidin-biotin-complex reagent for 1h at RT and developed with diaminobenzidine. Sections were mounted, dehydrated, and cleared in a series of alcohol washes and histoclear, then coverslipped. Number of PV cells in prelimbic cortex, dentate gyrus, CA1, and CA3 regions of the hippocampus was counted at 100× by two independent raters. Inter-rater reliability was 95% and raters were blind to treatment group.

### 2.6. Cannulae and Hippocampal Drug Infusions

See supplement. Briefly, rats were anesthetized, and bilateral guide cannulae were implanted into dorsal hippocampus. Animals were allowed to recover for 5d.

Drugs were infused for 2min at a rate of 0.25μl/min/side via infusion cannulae (1.5mm projection) using a microliter syringe and pump (KD Scientific). Infusers were kept in place for 2min after infusion was complete (Burnham et al. 2016).

### 2.7. Experimental Designs

(see Fig 1A)

**Fig 1.**
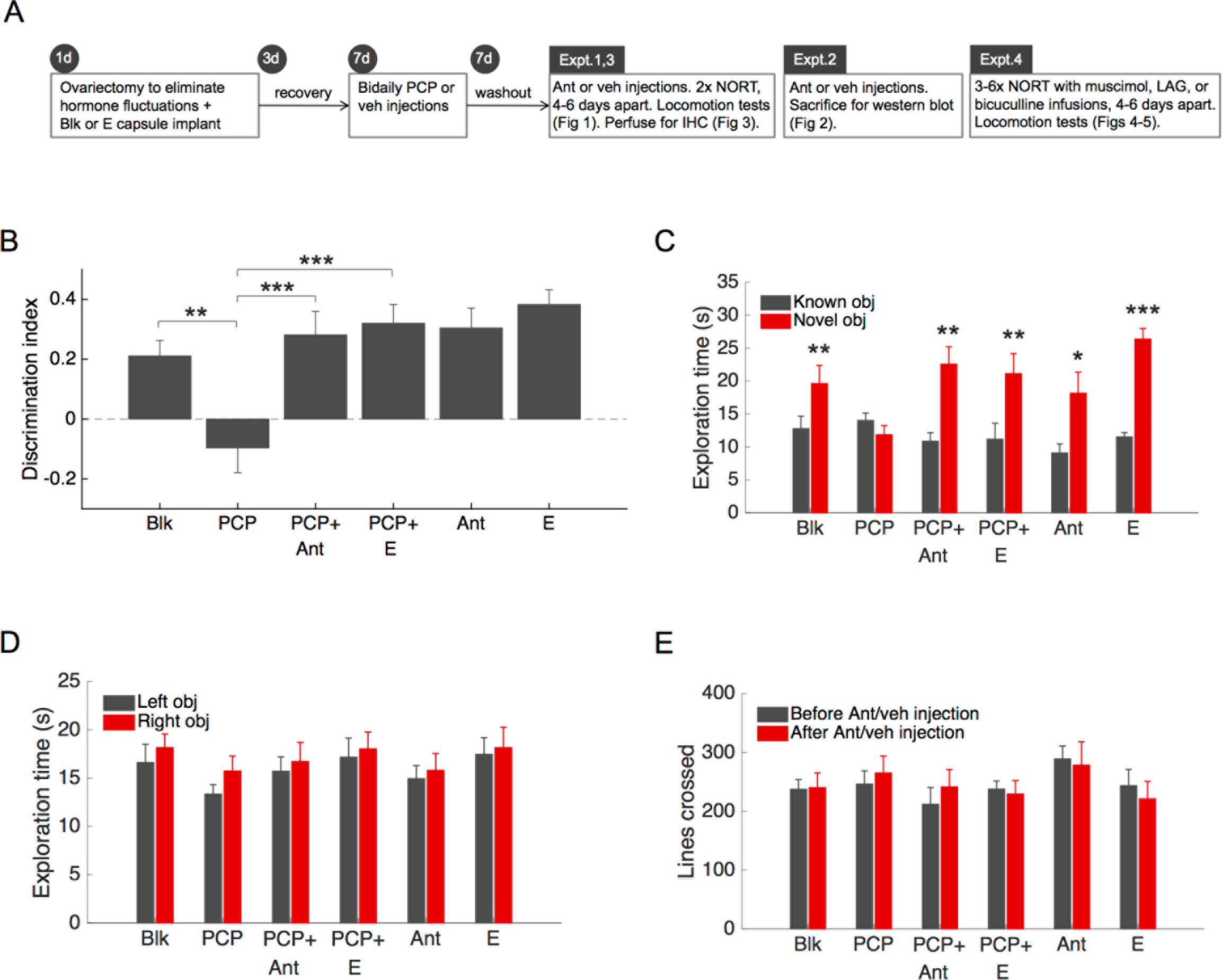
Suppression of LH with Antide or estradiol rescued recognition memory in ovx PCP-treated females. A. Experimental schedule B. Performance in novel object recognition task NORT). Sub-chronic PCP disrupted object discrimination compared to vehicle (PCP vs Blk). Blocking LH with Antide reversed PCP—s effects (PCP+Ant vs PCP), as did estradiol (PCP- E). Neither Ant nor E significantly affected object discrimination relative to Blk controls. Zero represents chance performance. C. Exploration time for novel vs. known objects during test trial. Consistent with discrimination index analysis, each group showed preference for the novel object, with the exception of the PCP group. D. There were no significant differences in total object exploration or left-right object preference amongst any of the groups during NORT exposure trial. E. There were no significant differences in number of lines crossed during locomotion tests, either across groups or before and after drug treatments. n = 7-10 rats/group. 2 tests/rat. Asterisks indicate statistically significant differences (* = p < .05, ** = p < .01, *** = p <.001).

#### 2.7.1 Expt 1. Effect of estradiol and Antide on PCP-induced memory loss

Rats were ovariectomized, implanted with Blk or E capsules, then treated with the PCP or vehicle (veh) regimen and allowed to recover. Rats were injected with Antide or veh 6h before each NORT. The six treatment groups were: blank implant+veh+veh (Blk; n=10); blank implant+PCP+veh (PCP: n=9); blank implant+PCP+Antide (PCP+Ant: n=10); E implant+PCP+veh (PCP+E: n=10). As additional controls, other females were blank implant+veh+Antide (Ant: n=7); E implant+veh+veh (E: n=10). Females were tested twice, with 4-6d between tests. Locomotion tests occurred before and after Antide/veh administration.

#### 2.7.2. Expt 2. Effect of PCP, estradiol, and Antide on GAD67 expression

Rats were treated as in Expt. 1. In place of the first NORT, rats were sacrificed for western blot procedure above. Final group numbers for dorsal hippocampal western blots were Blk n=11, PCP n=10, PCP+Ant n=7, PCP+E n=12, Ant n=7, E n=9. Final group numbers for prelimbic western blots were Blk n=9, PCP n=10, PCP+Ant n=11, PCP+E n=11, Ant n=8, E n=9.

#### 2.7.3. Expt 3. Effect of PCP, estradiol, and Antide on parvalbumin-expressing cells

Immediately after the second NORT, rats from Expt. 1 were anesthetized, perfused, and brains were analyzed using the immunohistochemistry and cell-counting protocol above. Final group numbers were Blk n=10, PCP n=8, PCP+Ant n=10, PCP+E n=8, Ant n=8, E n=8.

#### 2.7.4. Expt 4. Effects of hippocampal infusion of GABAergic drugs on NORT

Rats were ovariectomized, implanted with Blk or E capsules, fitted with guide cannulae, and treated with PCP or vehicle. During PCP washout, animals were habituated to the infusion procedure (see supplement). For behavioral testing, each female was infused with drug or vehicle (0.5μl sterile saline/side) 20-25min before NORT.

Expt 4A. To determine if the GABA_A_ agonist muscimol reverses NORT deficits induced by PCP, ovx, PCP-treated females (n=10) received muscimol (200ng/0.5μL/side; Sigma Aldrich, St. Louis, MO) or vehicle prior to NORT. Animals were infused in a repeated-measures counterbalanced design with three NORT tests under both control and muscimol conditions, with 4-6d between tests.

To assess locomotion animals received three locomotion tests: the day before the first NORT (no infusion), immediately after the second NORT (infusion of first drug), and immediately after the fifth NORT (infusion of second drug).

Expt 4B. To determine if the GABA_A_ antagonist bicuculline methiodide (bicuculline) blocks NORT performance in ovx females, similar to the effect of PCP, Blk animals (n=8) received either bicuculline (250ng/0.5μl/side; Sigma Aldrich, St. Louis, MO) or vehicle in a repeated-measures, counterbalanced design. Drug administration parameters and behavioral testing were identical to Expt. 4A.

Expt 4C. To determine if decreasing GABA synthesis also blocks NORT performance in ovx females, Blk females (n=6) received either L-allylglycine (LAG; 20μg/0.5μl/side), or vehicle in a repeated-measures, counterbalanced design identical to Expt. 4A.

Expt 4D. To determine if the GABA_A_ antagonist bicuculline prevents the ability of E to rescue NORT performance deficits caused by PCP, PCP+E females received infusions of either bicuculline (n=6, 250ng/0.5μl/side) or vehicle (n=6) in an independent groups design. Females underwent 3 tests with 4-6d between tests. Locomotion tests occurred immediately after the second NORT.

After testing, animals from Expts 4A-D were anesthetized, decapitated, and cannulae placement was verified using cresyl violet staining (see supplement).

### 2.8. Statistical Analysis

Data are expressed as mean ± SEM. NORT data, total object exploration, western blot, and PV cell counts were analyzed using one-way ANOVA. Data from each female were averaged prior to determination of group means. Locomotion tests were analyzed using two-way repeated-measures ANOVA, with treatment group and test number as the two factors. NORT data for drug-infused animals and left-right object preference were analyzed using within-subjects t-tests. Pairwise t-tests with pooled standard deviation were used to compare groups of pre-determined interest after ANOVA (Blk vs. each other treatment group, PCP vs. PCP+Ant, and PCP vs. PCP+E), and the Benjamini-Hochberg false-discovery rate method was used to correct for multiple comparisons (Benjamini and Hochberg 1995). An alpha of .05 was used as the threshold for statistical significance. Data were verified to fit the assumptions of statistical tests before their use (see supplement S8-S12). All statistical tests were performed using R.

## 3. RESULTS

### 3.1. Expt 1. Estradiol and Antide mitigate PCP-induced memory loss

Subchronic PCP treatment caused loss of recognition memory (Fig 1), which was rescued by inhibition of LH with Antide (a GnRH antagonist) or treatment with estradiol. There was a significant effect of treatment on discrimination index in NORT (F_5,50_=6.54, p=.000095, Fig 1B). Subchronic PCP caused deficits of object recognition memory relative to Blk controls (PCP vs. Blk p=.002). Antide rescued recognition memory in PCP-treated animals (PCP+Ant vs. PCP p=.0002; PCP+Ant vs. Blk p=.44). Estradiol replacement after ovariectomy prevented PCP-induced loss of recognition memory (PCP+E vs. PCP p=.00004; PCP+E vs. Blk p=.23). Estradiol and Antide had no significant effect on recognition memory in non-PCP-treated control animals (E vs. Blk p=.063; Ant vs. Blk p=.35, see supplement S1 for details). Consistent with this, each treatment group other than the PCP group showed strong novel object recognition, as evidenced by the significantly greater amount of time spent exploring the novel than the familiar object (Fig 1C,: PCP p=.30, all other groups p<.012).

Performance in NORT was not attributable to differences in exposure to objects during the exploration trial (Fig 1D). There was no significant effect of treatment on total object exploration during the exposure trial (F_5,50_=.80, p=.55). Moreover, no left-right object preferences within groups were found (Blk p=.45, PCP p=.18, PCP+Ant p=.60, Ant p=.68, PCP+E p=.46, E p=.77).

Differences in NORT performance were not caused by differences in locomotion (Fig 1E): There was no significant main effect or interaction between group treatment or drug injection/test number on lines crossed during locomotion testing (treatment F_5,50_=.86, p=.51; test number F_1,50_=0.00, p=.98; treatment-test number F5,50=.69, p=.64).

### 3.2. Expt 2. Effect of PCP, estradiol, and Antide on GAD67 expression

In the dorsal hippocampus, PCP decreased GAD67 expression (Fig 2A). Antide reversed this effect of PCP, restoring GAD67 to baseline expression, whereas estradiol failed to restore GAD67 in PCP-treated females. That is, western blots showed a significant difference between treatment groups in GAD67 expression in the dorsal hippocampus (one-way ANOVA: F5,50=3.3, p=.012). Subsequent analysis indicated that subchronic PCP treatment caused a ~20% reduction of GAD67 in dorsal hippocampus (PCP vs. Blk p=.014, see supplement S2 for details). Antide restored hippocampal GAD67 to control levels in PCP-treated animals (PCP+Ant vs. PCP p=.0071; PCP+Ant vs. Blk p=.44). PCP animals implanted with estradiol still showed a ~20% GAD67 deficit (PCP+E vs. PCP p=.59; PCP+E vs. Blk p=.036). In non-PCP-treated controls, neither Antide nor estradiol caused statistically significant differences in hippocampal GAD67 (Ant vs. Blk p=.68, E vs. Blk p=.81). There were no significant differences in β-actin expression across treatment groups (one-way ANOVA: F_5,50_=.28, p=.92; Fig S2A).

**Fig 2.**
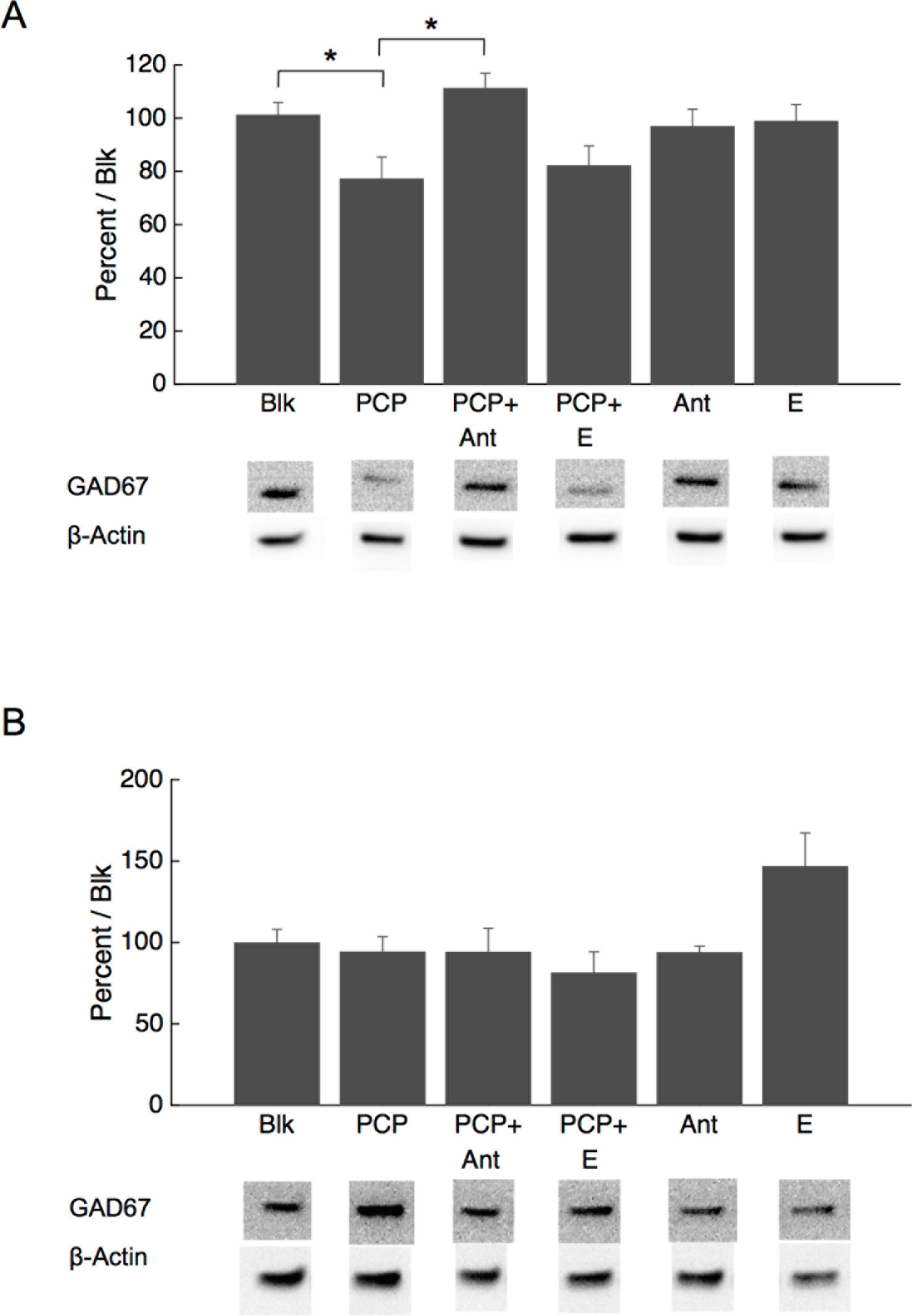
PCP decreased GAD67 expression in dorsal hippocampus and inhibition of LH restored it. A. GAD67 expression in dorsal hippocampus, measured by western blot. Hippocampal GAD67 was significantly decreased in PCP-treated rats relative to controls (PCP vs Blk). Inhibition of LH (PCP+ Ant) significantly increased hippocampal GAD67 (PCP+Ant vs PCP). B. GAD67 expression in prelimbic cortex. ANOVA revealed a significant difference in GAD67 expression, however post hoc tests indicated no effect of PCP (PCP vs Blk) or any other treatment. All GAD67 data were normalized over P-Actin and intra-gel blank controls. Representative blots are shown below. Original images for representative blots are shown in supplementary **Fig S1**. β-actin expression, shown in **Fig S2**, did not differ across treatment groups. n = 7-12 rats/group. Three technical replicates were performed for hippocampal data. Asterisks indicate statistically significant differences (* = p < .05).

In the prelimbic cortex (Fig 2B) there was no clear effect of PCP, estradiol, or Antide on GAD67 expression. Western blotting revealed a significant effect of treatment on GAD67 expression in prelimbic cortex (F_5,50_=3.3, p=.012). However post-hoc tests revealed no significant differences between PCP and Blk (Blk vs. PCP p=.77, see supplement S3 for details). There were no significant differences in β-actin expression across treatment groups (one-way ANOVA: F_5,50_=.17, p=.97; Fig S2B).

### 3.3. Expt 3. Effect of PCP, estradiol, and Antide on the number of PV immunoreactive cells

These treatments did not clearly affect the number of PV expressing cells in the brain regions examined (Fig 3). No significant differences in number of PV-immunoreactive cells were found in either the prelimbic cortex (Fig 3A, F_5,47_=1.29, p=.28), dentate gyrus (Fig 3B, F_5,34_=1.1, p=.38), or CA3 region of the hippocampus (Fig 3C, F_5,35_=2.37, p=.059). Although there was a significant effect of treatment on PV-expressing cell number in CA1 of the hippocampus (Fig 3D, F_5,35_=2.51, p=.048), PCP did not differ significantly from control, and post-hoc tests were unable to localize any statistically significant differences (see supplement S4 for p-values).

**Fig 3.**
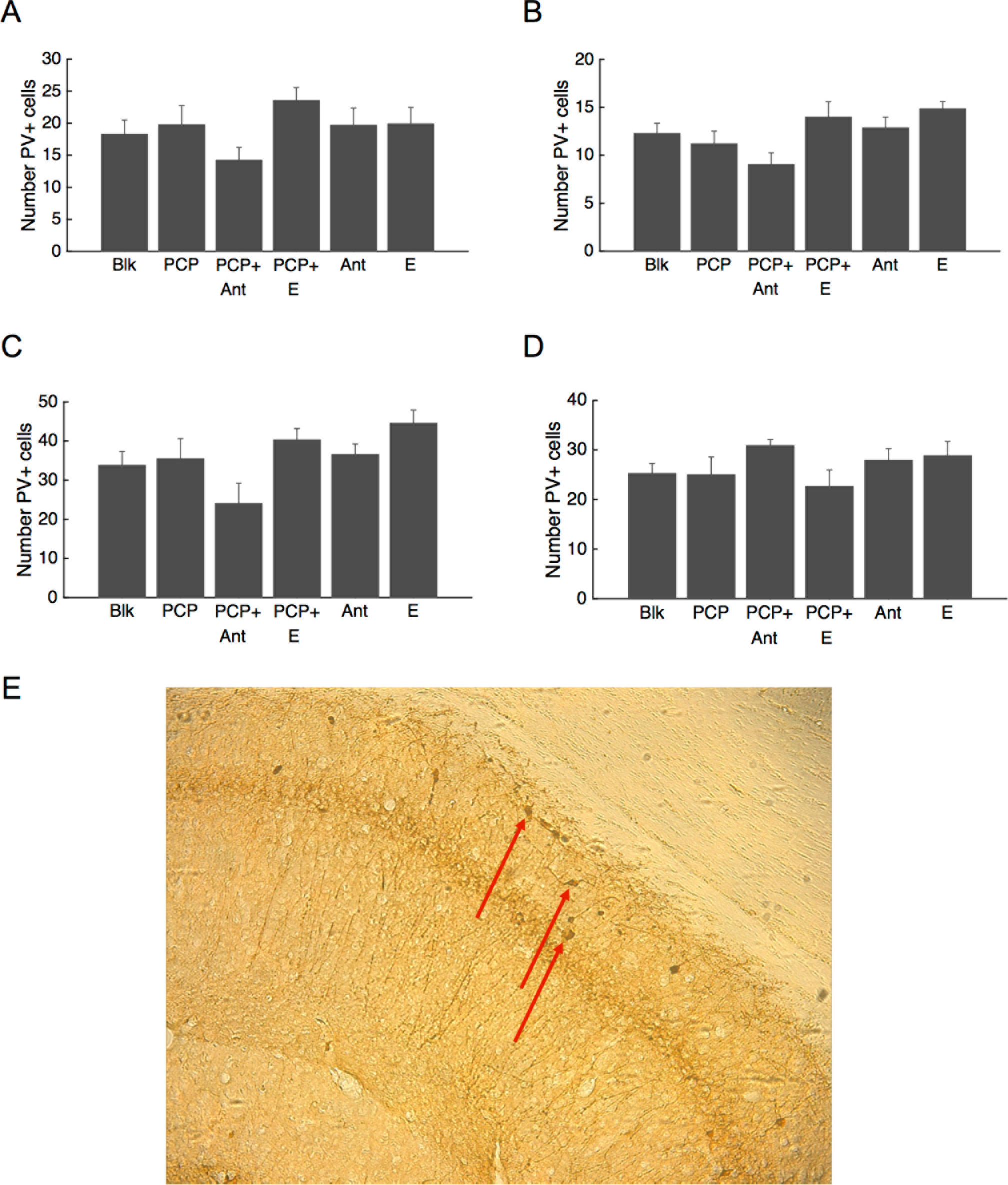
Number of immunoreactive parvalbumin-expressing (PV+) cells was unaffected by PCP and hormonal treatments (n = 8-10). ANOVA revealed no significant differences in PV cell number in the prelimbic cortex (panel A), dentate gyrus (panel B), or CA3 region of the hippocampus (panel C). ANOVA revealed a significant difference in cell counts in the CA1 region of the hippocampus (p = .048), however this effect was not localizable with post hoc tests (D) and PCP did not differ significantly from control PCP vs Blk. E. Representative immunohistochemical staining against PV. Arrows point to several example cells.

### 3.4. Expt 4. Stimulation of GABA receptors in the hippocampus reverses PCP-induced memory loss, and blocking hippocampal GABA receptors or synthesis causes memory loss

Infusion of the GABA_A_ receptor agonist muscimol into the dorsal hippocampus counteracted the amnesic effects of PCP (Fig 4A-B). Consistent with Expt 1, PCP-treated animals did not show novel object recognition, as there was no clear differential exploration of the novel object (p=.50). Infusion of GABA_A_ receptor agonist muscimol into dorsal hippocampus rescued recognition memory in PCP-treated ovx females, as evidenced by both the discrimination index (p=.010) and difference in exploration times of the known and novel objects (p=.0044). Differences in NORT performance were not due to differences in object exploration during exposure trial (t-test p=.53), and no left-right object preferences were found (PCP+muscimol p=.49, PCP+veh p=.61, Fig S3A). Moreover, there was no significant main effect or interaction of treatment or test number on lines crossed in locomotion tests (Fig S4A, and see supplement S5).

**Fig 4.**
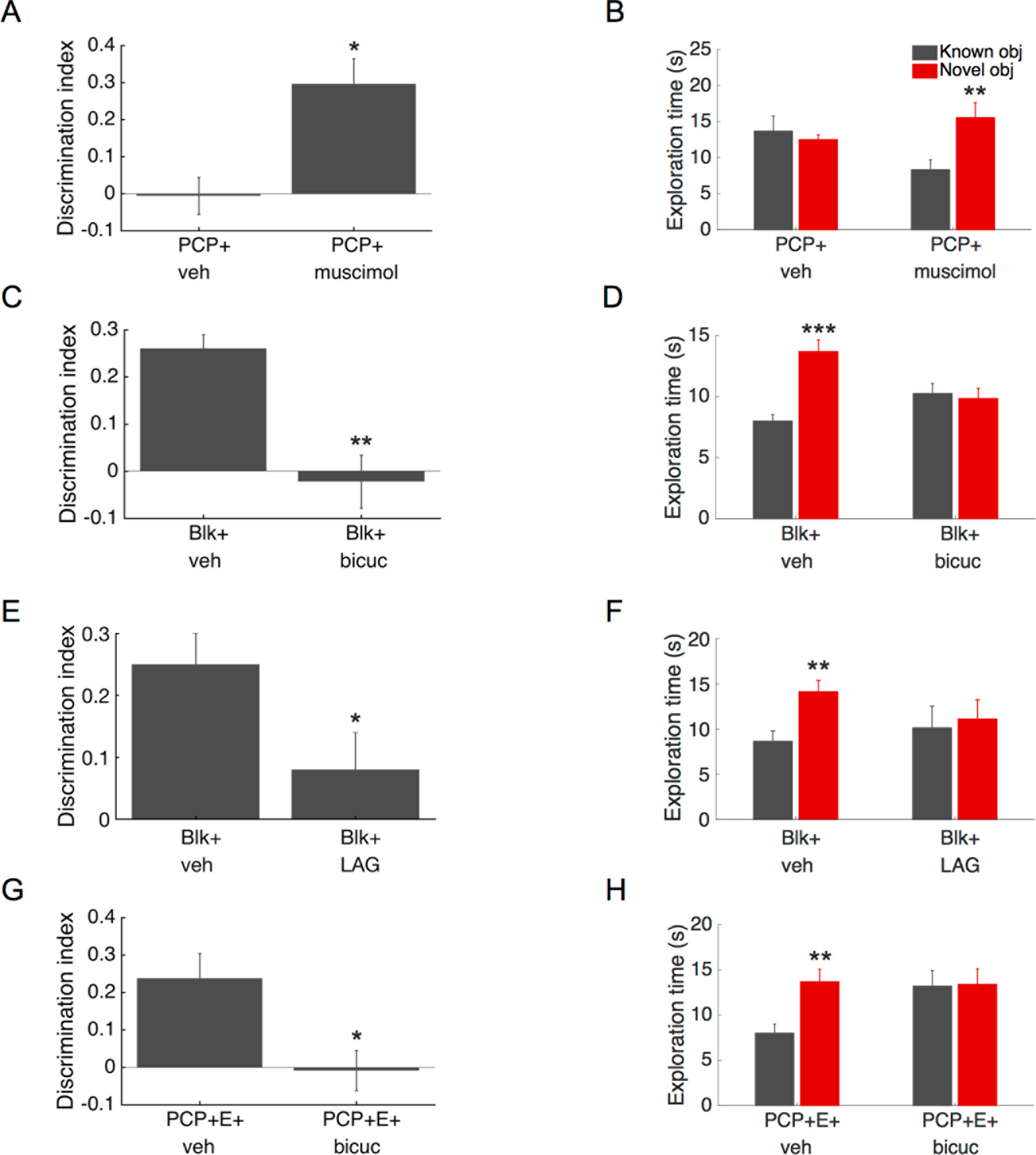
Stimulating hippocampal GABA_A_ receptors rescued recognition memory in PCP-treated rats. Blocking hippocampal GABA_A_ receptors or GABA synthesis impaired recognition memory in Blk and estrogen-PCP-treated animals. A. Infusions of the GABA_A_ agonist muscimol into dorsal hippocampus significantly raised novel object discrimination in PCP-treated rats relative to vehicle infusion controls (n=10, 3 tests/drug condition). B. Consistent with discrimination index, PCP animals showed chance-level exploration of novel vs. known objects after vehicle infusion and significantly greater exploration of novel objects after muscimol infusion. C. Infusions of the GABA_A_ antagonist bicuculline into dorsal hippocampus impaired NORT performance in non-PCP-treated Blk rats (n=8, 3 tests/drug condition). D. Data from C expressed as exploration time (Blk+veh, Blk+bicuc). E. Similarly, infusions of the GAD inhibitor LAG into dorsal hippocampus impaired NORT performance in non-PCP-treated Blk rats (n=8, 3 tests/drug condition). F. Data from E expressed as exploration time (Blk+veh, Blk+bicuc). G. Infusions of GABA_A_ antagonist bicuculline into dorsal hippocampus disrupted NORT performance in rats treated with both PCP and estradiol (n=6, 3 tests/animal). H. Data from G expressed as exploration time. Asterisks indicate statistically significant differences compared to veh (A, C, E) or to known object in same treatment group (B, D, F). (* = p < .05, ** = p < .01, *** = p <.001).

Blocking GABA_A_ receptors in the dorsal hippocampus caused memory loss that was comparable to that observed following PCP administration (Fig 4C-D). Infusion of GABA_A_ receptor antagonist bicuculline into the dorsal hippocampus of control (non-PCP treated) females caused loss of recognition memory (discrimination index p=.0011; known vs. novel: Blk+veh p=.00016, Blk+bicuculline p=.72). Differences in NORT performance were not due to differences in exploration during the exposure trial (p=.36, Fig S3B), left-right preferences (Blk+bicuculline p=.55, Blk+veh p=.65), or locomotion differences (Fig S4B, see supplement S6).

Consistent with the effect of GABA_A_ receptor antagonism, blocking GABA synthesis in the dorsal hippocampus also caused memory loss. Infusion of the GAD inhibitor LAG into the dorsal hippocampus of control (non-PCP treated) females caused loss of recognition memory (discrimination index p=.022; known vs novel: veh p=.0027, LAG p=.36). There were no significant differences in exploration during the exposure trial (p=.12, Fig S3C) or left-right preference (LAG p=.95, veh p=.40). Although there was a significant effect of treatment on locomotion, post hoc analysis found no differences in locomotion following administration of LAG vs. vehicle (p=.41, Fig S4C). The only significant difference found was between testing before treatment and testing after administration of vehicle (p=.018).

Blocking GABA_A_ receptors in dorsal hippocampus also eliminated estradiol’s rescue of memory in PCP-treated females (Fig 4E-F). Dorsal hippocampal infusion of bicuculline impaired novel object discrimination in PCP-treated animals that were implanted with estradiol (discrimination index p=.016; known vs. novel: E+PCP+veh p=.009, E+PCP+bicuculline p=.90). There were no differences in object exploration during the exposure trial (t-test p=.11, Fig S3D), left-right preferences within groups (E+PCP+bicuculline p=.76, E+PCP+veh p=.18), or locomotion differences (p=.46, Fig S4D).

Cresyl violet histology verified that infusion cannulae were appropriately placed in dorsal hippocampus (Fig 5).

**Fig 5.**
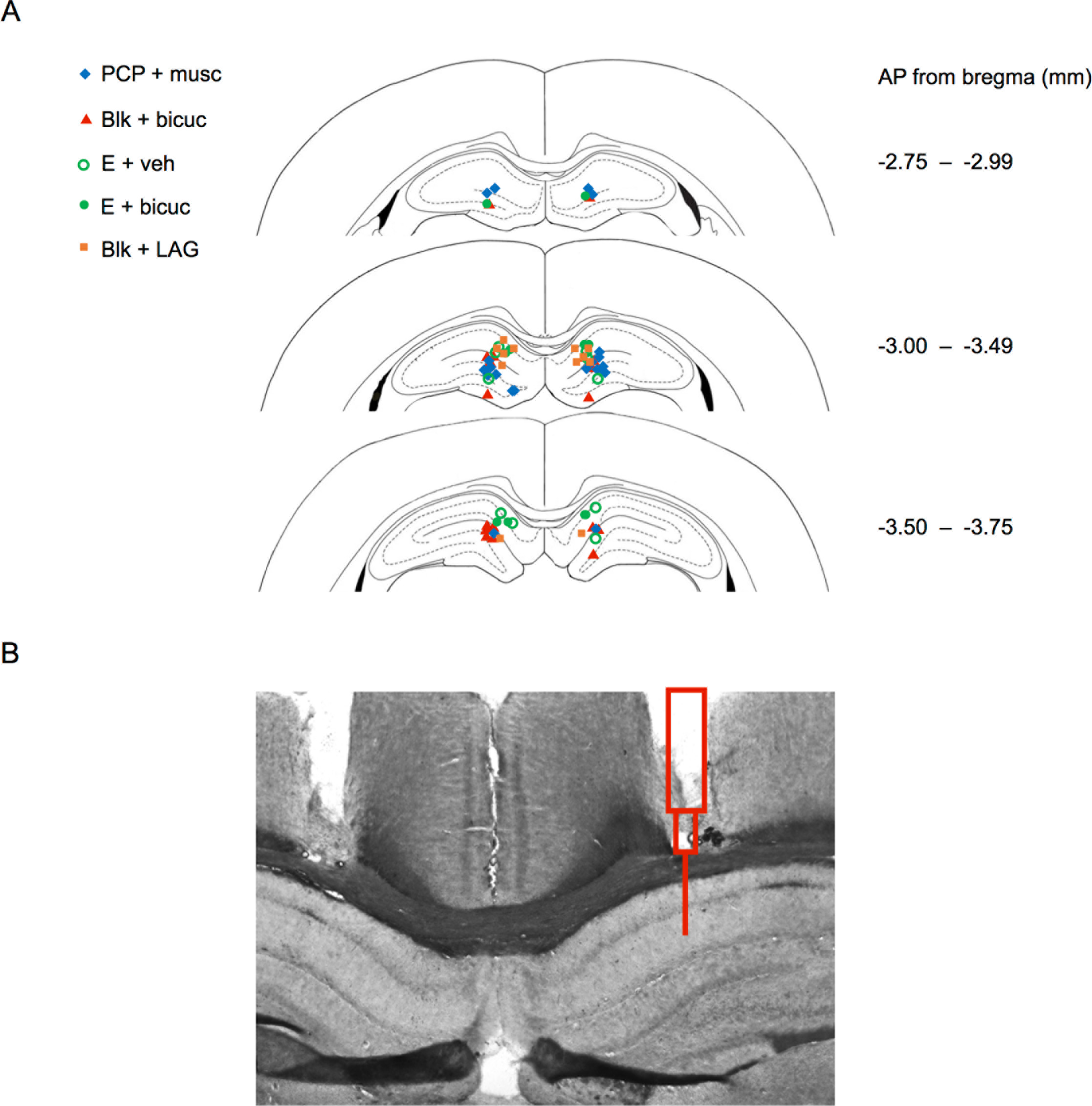
Location of infusion cannulae in dorsal hippocampus. A. Dots show final location of bilateral infusion cannulae in PCP, Blk, and PCP+E animals. B. Representative cresyl-violet-stained brain section used to verify cannulae placement, with schematic of cannula superimposed over one hemisphere.

## 4. DISCUSSION

Here we demonstrate that modulation of LH and estradiol can rescue recognition memory in an NMDAR antagonist model of schizophrenia. We found that PCP decreased expression of hippocampal GAD67, an effect that was reversed by inhibition of LH, but not administration of E. Localized hippocampal infusion of the GABA agonist muscimol reversed NORT deficits induced by PCP. Consistent with this, the GABA antagonist bicuculine or GAD-inhibitor LAG decreased NORT in ovx females and blocked enhancement of NORT exerted by E in PCP-treated females. This suggests that GABAergic changes may contribute both to PCP’s disruption of recognition memory and the hormones’ ability to counteract this effect. Importantly, the observed effects are not due to changes in locomotion or object exploration during the exposure trial.

These findings are consistent with the idea that PCP-induced amnesia is mediated by GABAergic disruptions specific to dorsal hippocampus. PCP reduced expression of hippocampal GAD67, which presumably decreases the amount of GABA available for release when inhibitory neurons fire (Lewis et al. 2012). Previous studies suggest that a reduction of GAD67 can upset the excitatory-inhibitory balance in dorsal hippocampus, which is thought to be an important component of short-term memory (Lewis et al. 2012; Lisman 2010; Sigurdsson et al. 2010). Our data then suggest a causal link between this localized GABAergic deficit and PCP-induced loss of recognition memory. Because dorsal hippocampal infusion of a GABA_A_ agonist rescued recognition memory in PCP-treated females and infusion of a GABA_A_ antagonist and GABA synthesis blocker produced memory loss similar to that of PCP, our data are consistent with the notion that disrupted GABA transmission in the hippocampus is both necessary and sufficient for PCP’s amnesic effects in this animal model. These findings complement previous work showing that complete lesions of dorsal hippocampus cause loss of object recognition memory in rodents, and support the idea that aberrant signals from the hippocampus may be capable of disrupting forms of object recognition typically thought to reside in regions such as perirhinal cortex (Broadbent et al. 2004; Cohen et al. 2013; Oliveira et al. 2010). Caution with the interpretation that a reduction in GAD67 underlies the memory deficit is warranted, however, as we did not directly rescue hippocampal GAD67 in PCP-treated animals. It is possible that our muscimol infusions stimulated GABA_A_ receptors to a greater extent than rescuing GAD67 would have (Hafting et al. 2008). As such the present results imply localization of PCP’s effects to dorsal hippocampus but do not provide direct evidence that decreased GAD67 causes the memory loss (Hafting et al. 2008).

These data suggest that the prefrontal cortex does not play a role in the effects of PCP on object recognition memory. This notion is supported by studies showing that lesioning prefrontal cortex does not affect performance in NORT and related tasks (Barker and Warburton 2011). Although other investigations have found that NMDAR antagonists cause loss of GAD67 specific to PV neurons in prelimbic cortex, these studies did not measure overall regional expression of GAD67 (Amitai et al. 2012; Behrens et al. 2007; Kinney et al. 2006). Because PV-expressing neurons are estimated to make up only 30-40% of the inhibitory neurons in rat frontal cortex, it is possible that PV-neuron-specific deficits of GAD67 were masked in our study by unaffected GAD67 within other GABAergic neuronal subtypes (Xu et al. 2010).

We did not observe a change in the expression of PV, a result that is inconsistent with some and consistent with other previous research. There is a current conflict as to whether NMDAR antagonists cause a long-term reduction of PV-expressing cells in cortex and hippocampus (Abdul-Monim et al. 2007; Behrens et al. 2007; Benneyworth et al. 2011; Rujescu et al. 2006; Turner et al. 2010; Wang et al. 2008). Such inconsistencies suggest that PCP’s effect on PV expression may be variable across different rat strains, sexes, or PCP doses. Overall, this lack of observed differences in PV cell counts suggests that PCP does not disrupt recognition memory by eliminating PV-expressing cells.

The present results suggest that either increasing E or decreasing LH may counteract loss of recognition memory in schizophrenia, supporting the idea that E’s therapeutic mechanisms include suppression of LH. PCP-induced memory loss occurred only in animals with high LH (i.e., Blk group), and inhibiting LH reversed this amnesic effect. As in previous studies, estradiol-treated animals, that also exhibit low LH, were invulnerable to PCP’s effects in the present paradigm (Roseman et al. 2012; Sutcliffe et al. 2008). As LH and its analogues are capable of disrupting memory on their own (Burnham et al. 2016; Burnham and Thornton 2015; Ziegler and Thornton 2010), our data indicate that both LH and PCP can impair memory. The marked increase in incidence of schizophrenia at menopause (McGregor et al. 2017) may be explained by such a combined mechanism, whereby high levels of LH interact with genetic or environmental factors in vulnerable individuals, leading to an increased probability in the onset of schizophrenia. Given that Antide was able to rescue recognition memory long after PCP washout, our data suggest that inhibiting LH might be able to restore some forms of memory in such affected individuals.

These data indicate that LH may affect hippocampal GABA neurotransmission, a relationship that has not been previously characterized. Inhibition of LH with Antide restored hippocampal GAD67 in PCP-treated animals to baseline control levels. LH has been found in the cytoplasm of hippocampal neurons in both rats and humans, and receptors for LH are found on neurons in the same region (Burnham and Thornton 2015). Currently, the mechanism through which LH affects GABA is unknown. One speculative possibility is that binding of LH to hippocampal LH receptors causes second messenger cascades that downregulate production of GAD67. This anti-GABAergic action could be responsible, at least in part, for the LH-induced cognitive impairment observed in this and other contexts (Burnham et al. 2016; Burnham and Thornton 2015; Casadesus et al. 2007; Ziegler and Thornton 2010).

Interestingly, estradiol reversed PCP’s amnesic effects but did not restore GAD67 expression in the hippocampus, leaving its mechanisms of memory rescue unknown. This is surprising, because estradiol inhibits release of LH and should have similar effects as Antide (Charlton 2008; Wise and Ratner 1980). However it is known that under constant E, animals still exhibit daily LH surges (Legan et al. 1975), indicating that E may be less effective than Antide at consistently suppressing LH (Ziegler and Thornton 2010). Still, GABA_A_ receptor inhibition blocked estradiol’s rescue of memory, supporting the notion that at least some of estradiol’s effects are dependent on the hippocampal GABA system. Estradiol has been shown to enhance GABA_A_ receptor binding in a number of brain regions, including the hippocampus, and hence may be reversing PCP’s effects by upregulating GABA receptors (Schumacher et al. 1989). Additionally, estradiol’s effects on hippocampal spine density or brain-derived neurotrophic factor could account for its ability to rescue memory, either through enhancement of GABA signaling (Ohba et al. 2005) or other mechanisms (Li et al. 2004; Wu et al. 2013).

These results may have clinical implications. Namely, our data provide conceptual support for models in which declarative memory loss in schizophrenia is caused by disrupted hippocampal inhibitory function (Lewis et al. 2012; Nakazawa et al. 2012). Hippocampal GABA abnormalities are reliably found in schizophrenia and other conditions such as bipolar disorder, and it is possible that selective GABAergic modulation could mitigate memory loss in such disorders (Benes et al. 2007; Curley et al. 2011; Lewis et al. 2012). These data also indicate that the hypothalamic-pituitary-gonadal axis may be an unexploited tool to aid in understanding mechanisms of cognitive loss in schizophrenia.

In conclusion, we demonstrate that inhibiting luteinizing hormone can rescue memory in an NMDAR-antagonist model of schizophrenia. Our data suggest that GABAergic loss specific to the dorsal hippocampus contributes to PCP- and LH-induced disruption of recognition memory.

## FUNDING AND DISCOLOSURES

Work was funded by Oberlin Grant-in-Aid and a Robert Rich Grant, from Oberlin College. The authors declare no conflicts of interest.

## ACKNOWLEDGEMENTS

We would like to thank Carey Lyons, Alexander Goldberg, Molly Martorella, and Hannah Rodgers for technical assistance.

---

**Fig S1.**
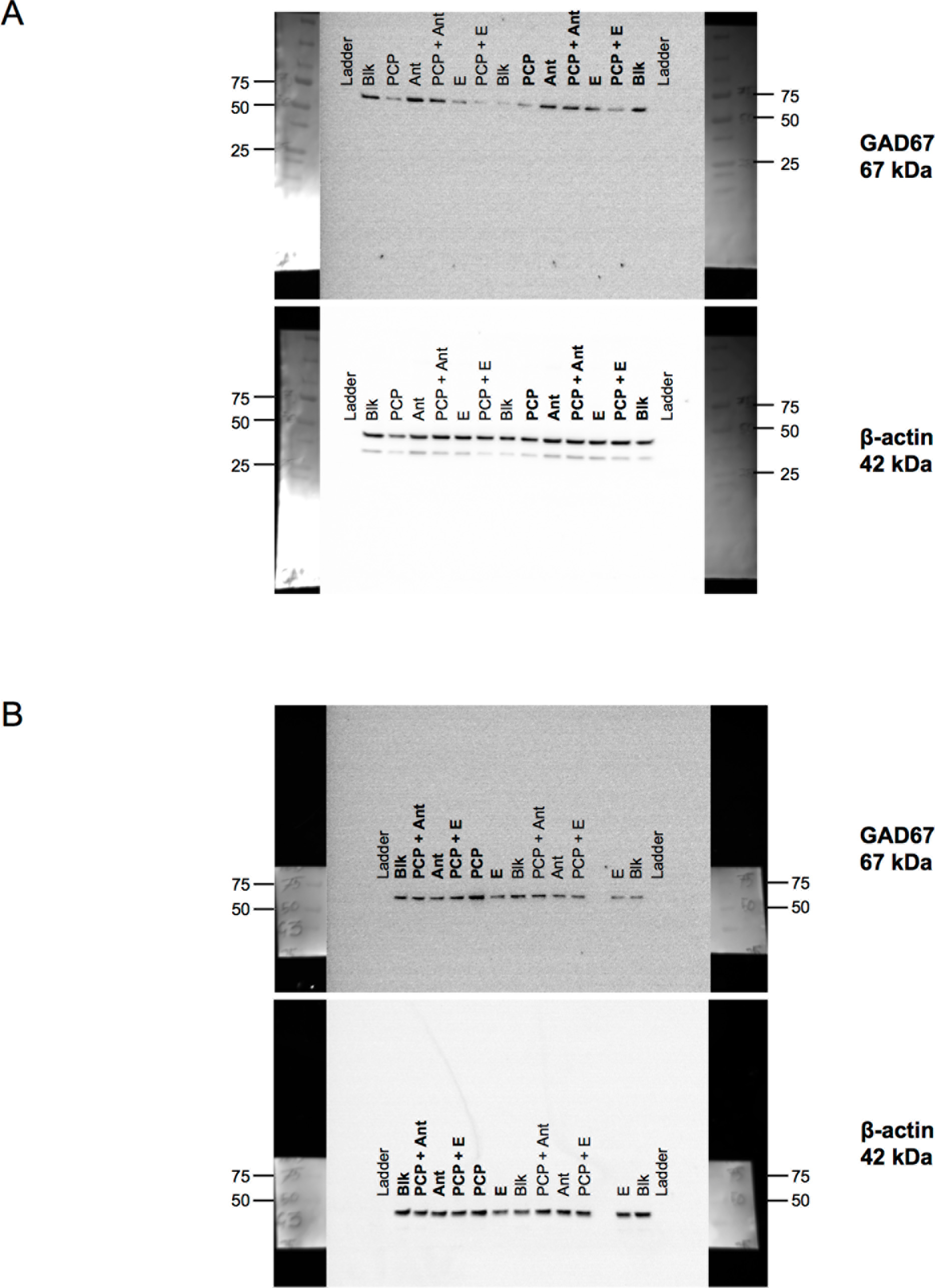
Original images of representative western blots from **Fig 2**. Treatment group is denoted above each band. Wells with bolded labels were rearranged for presentation in **Fig 2**. Left- and right-most wells of each blot were loaded with molecular-weight marker proteins (Ladder), which also served as negative controls. Blots are flanked by cropped photos of molecular weight markers. All blots appeared at the expected molecular weight for the proteins being analyzed (GAD67: 67kDa; β-actin: 42 kDa). A. Original images for hippocampus western blots in **Fig 2A**. B. Original images for prelimbic cortex western blots in **Fig 2B**.

**Fig S2.**
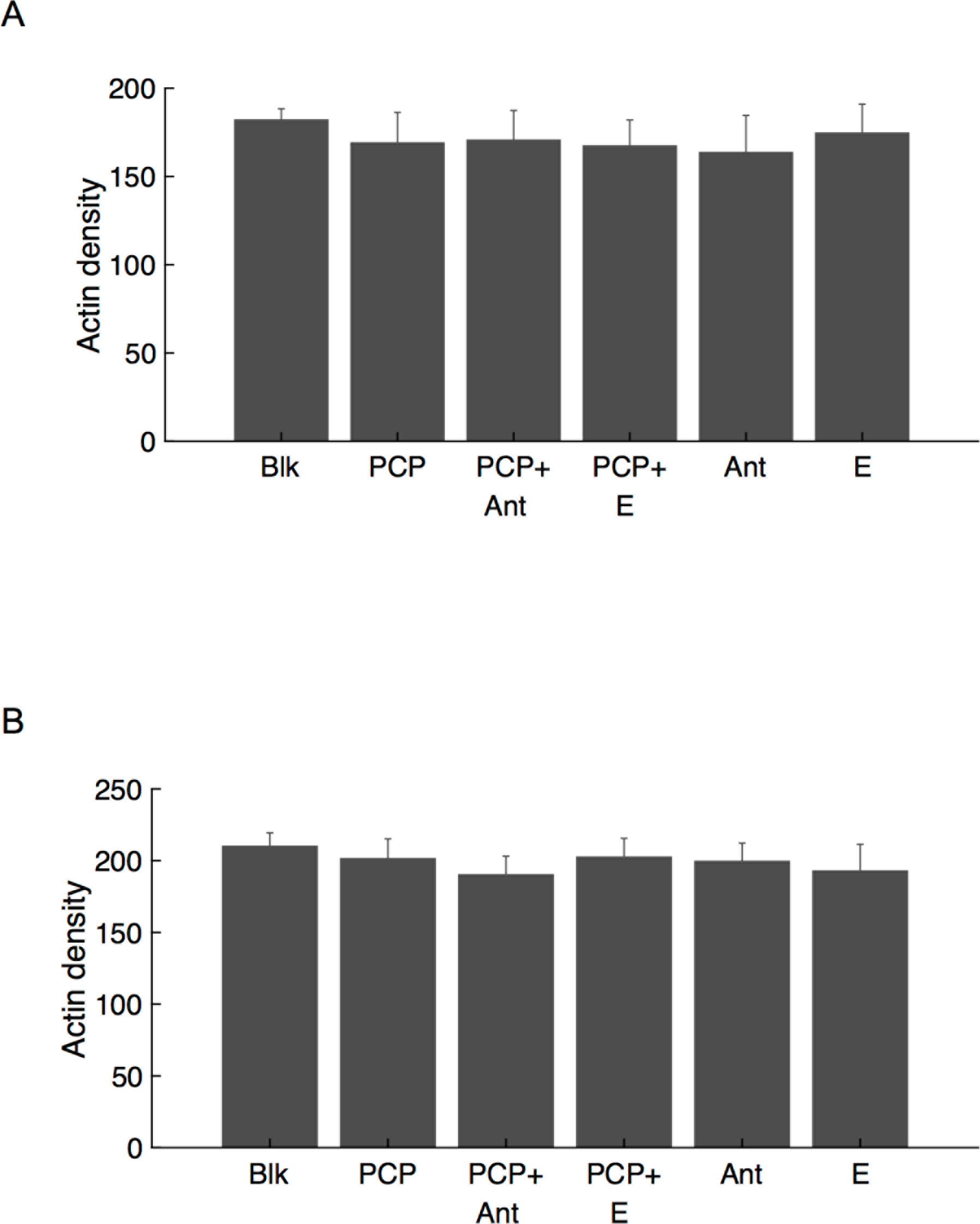
β-actin expression, used as a loading control for western blots in **Figs 2 and S1**, was unaffected by the drug and hormone treatments. That is, one-way ANOVA revealed no significant effect of treatment group on β-actin densitometry values in the dorsal hippocampus (A; F_5,50_=.28, p=.92) or prelimbic cortex (B; F_5,50_=.17, p=.97).

**Fig S3.**
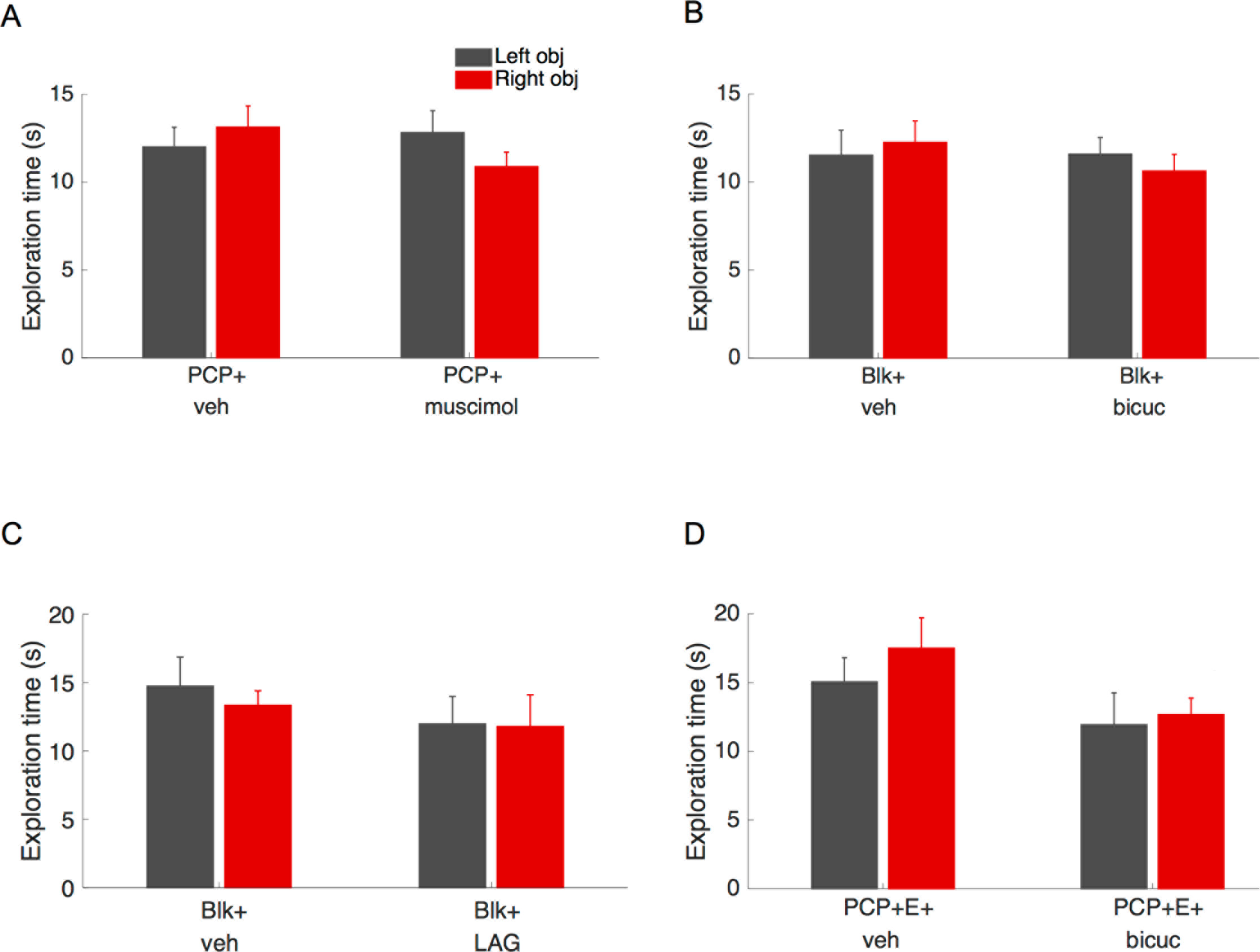
Dorsal hippocampal infusions did not alter object exploration during exposure trial of NORT. No statistically significant differences in total object exploration or left-right object preference were found in any of the experiments. A. PCP+muscimol infusion (Expt. 4A); B. Blk+bicuculline infusion (Expt. 4B); C. Blk+LAG infusion (Expt. 4C); D. PCP+E+bicuculline (Expt. 4D).

**Fig S4.**
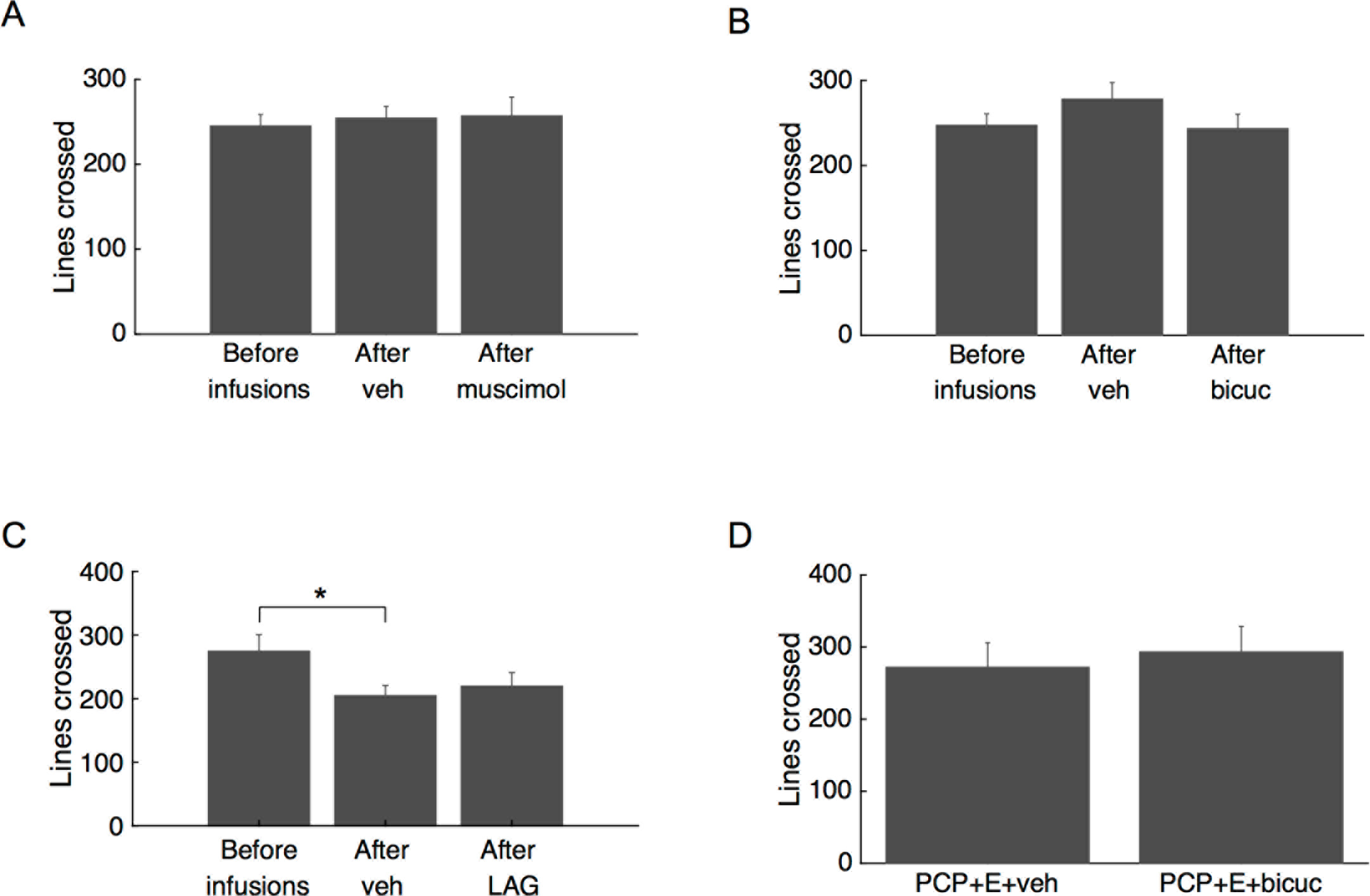
Dorsal hippocampal infusions of GABA antagonist/agonist did not alter the number of lines crossed in locomotion tests. There were no statistically significant differences in lines crossed among infused animals either before or after infusions of drug or vehicle. A. PCP/muscimol infusion experiment. No significant differences. B. Blk/bicuculline infusion expt. No significant differences before vs after drug treatment. The only statistically significant difference was between non-infused and vehicle-infused animals. C. Blk/LAG infusion expt. No significant differences. D. E+PCP/bicuculline infusion expt. No significant differences.

